# Connectome-harmonic decomposition of human brain activity reveals dynamical repertoire re-organization under LSD

**DOI:** 10.1101/163667

**Authors:** Selen Atasoy, Leor Roseman, Mendel Kaelen, Morten L. Kringelbach, Gustavo Deco, Robin L. Carhart-Harris

## Abstract

Recent studies have started to elucidate the effects of lysergic acid diethylamide (LSD) on the human brain but the underlying dynamics are not yet fully understood. Here we used ‘connectome-harmonic decomposition’, a novel method to investigate the dynamical changes in brain states. We found that LSD alters the energy and the power of individual harmonic brain states in a frequency-selective manner. Remarkably, this leads to an expansion of the repertoire of active brain states, suggestive of a general re-organization of brain dynamics given the non-random increase in co-activation across frequencies. Interestingly, the frequency distribution of the active repertoire of brain states under LSD closely follows power-laws indicating a re-organization of the dynamics at the edge of criticality. Beyond the present findings, these methods open up for a better understanding of the complex brain dynamics in health and disease.

## Introduction

The psychoactive effects of lysergic acid diethylamide (LSD) were discovered in 1943 and began to be fully reported on in the late 1940s^1^ and early 1950s^2^. For a period of approximately 15 years, LSD and related psychedelics were used as tools to explore and understand consciousness^3^, psychopathology^4^ and also to treat mental illness^5^. In addition to the LSD’s recognised effects on perception, it was reported to ‘loosen’ constraints on consciousness, expanding its breadth, without compromising wakefulness. Indeed, it was said of LSD and related psychedelics, that they could serve as ‘microscopes’ or ‘telescopes’ for psychology, allowing scientists to see more of the mind than is normally visible^6^. For various reasons^7^, psychology has been slow to embrace and exploit this powerful scientific tool, but now, with significant advances in modern functional neuroimaging and underperformance in medical psychiatry^8^, there are significant incentives to do so.

Previous functional magentic resonance imaging (fMRI) and magnetoencephalography (MEG) work has reported increased visual cortex blood flow, increased whole-brain functional integration (decreased modulatory organization), and decreased oscillatory power across a broad frequency range under LSD^9,10^. These findings are based on conventional neuroimage analysis methods and although they offer a useful impression of the neural correlates of the LSD state, they provide limited information on whole-brain dynamics. Recent research has begun to postulate changes in brain dynamics that may account for the putative broadening of consciousness under psychedelics^11^; however, experimental evidence for the hypothesized dynamical changes is limited^12^.

A fundamental limitation to understanding the nature of these spatio-temporal changes, is the lack of mathematical tools and methods particularly tailored to grasp the complex dynamics of cortical activity^13,14^. Here we describe a novel method to decompose resting-state fMRI data under LSD and placebo into a set of independent, frequency-specific brain states. The unique decomposition of cortical activity into a sum of frequency-specific brain states allows us to characterize not only the dominant frequency content of brain activity under LSD and placebo but also the underlying brain dynamics in terms of a frequency distribution among the complete repertoire of these states.

Our technique capitalizes on the identification of the brain states as harmonic modes of the brain’s structural connectivity and the decomposition of fMRI-measured cortical activity patterns into these harmonic brain states. The estimation of brain states builds on the previously described connectome-harmonic framework^15^ (Fig. 1), which was developed to explain how patterns of synchronous neural activity emerge from the particular structural connectivity of the human brain - the human connectome. The utilization of connectome harmonics as brain states composing complex cortical dynamics offers two important advantages; firstly, the connectome harmonics, by definition, are the spatial extension of the Fourier basis to the particular structural connectivity of the human brain, the human connectome^15^ (Fig. 1**a-f**, Fig. S1), enabling for the first time a spatial harmonic analysis tailored to the human connectome. Secondly, they correspond to spatial cortical patterns formed by fully synchronized neural activity, each associated with a different temporal frequency as well as a different spatial wavelength, as exemplified in the general phenomenon of standing waves and in cymatic patterns^15^. Hence, they provide a set of independent and fully synchronous brain states (cortical patterns) with increasing spatial frequency that is also particularly adapted to the anatomy of the human brain. Mutually, the activation of these harmonic brain states composes the complex spatio-temporal dynamics of cortical activity. Hence, the unique decomposition of the cortical activity patterns into the set of connectome harmonics identifies the individual contribution of each harmonic brain state (Fig. 1**g-h**). Importantly, this connectome-specific harmonic decomposition of cortical activity enables the evaluation of fundamental properties of these harmonic brain states such as energy and power, hence introducing these well-studied physical concepts as novel tools for neuroscience.

**Figure 1.**
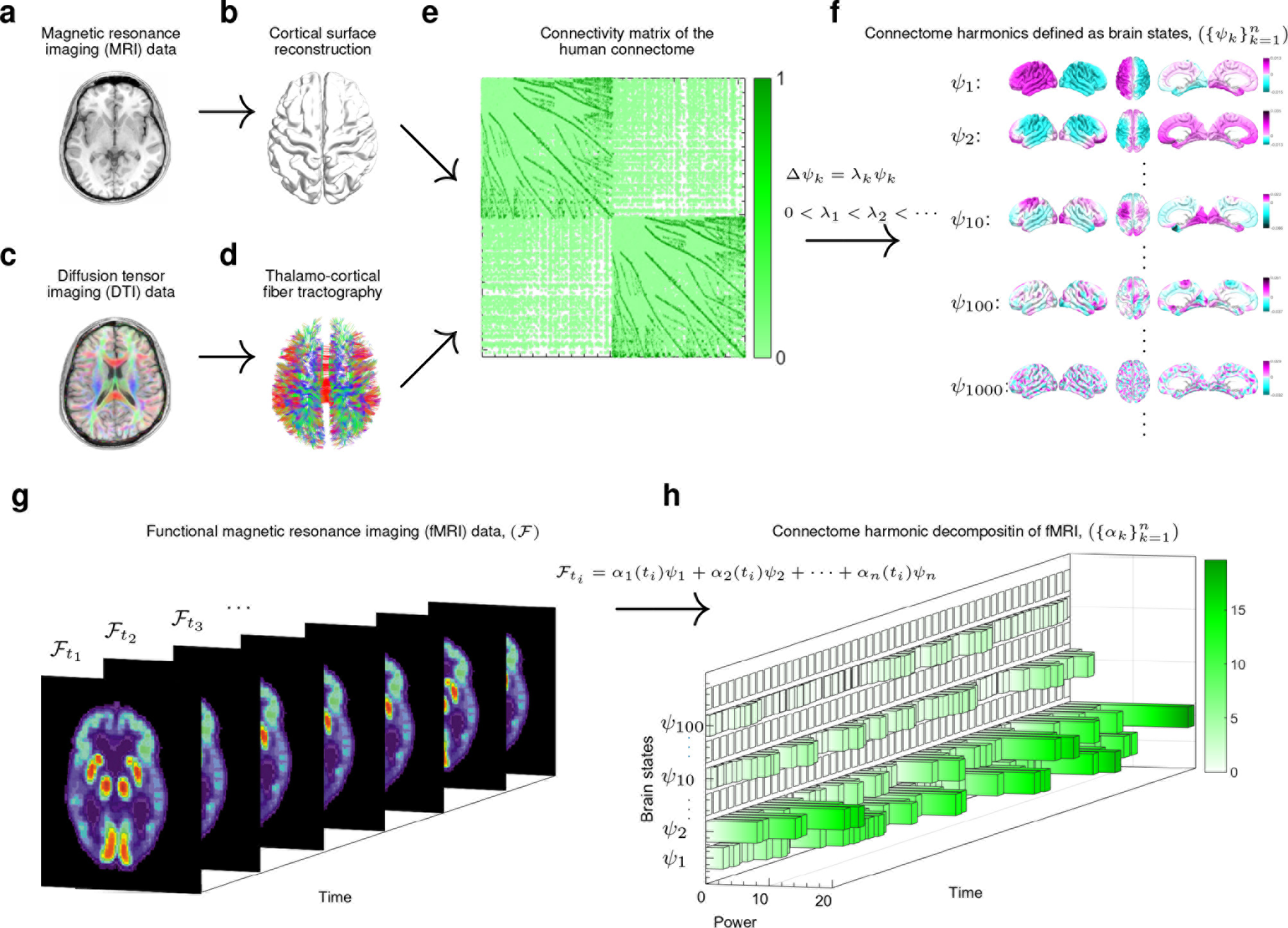
Illustration of the workflow. T1 magnetic resonance imaging (MRI) data (**a**) is used to reconstruct the cortical surface between gray and white matter as shown in (**b**). Diffusion tensor imaging (DTI) deta (**c**) is used to extract thalamo-cortical fibers using deterministic fiber tractography as shown in (**d**). Both, local and long-distance connections are combined to create the connectivity matrix of the human connectome as illustrated in (**e**)i Connectome harmonics 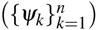 (**f**) are estimated by applying the Laplace operator Δ to human connectome and computing its eigenvectors 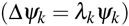 Functional magnetic resonance imaging (fMRI) data 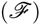 as illustrated in (**g**) is decomposed in to the activation of connectome harmonics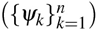 yielding the power ot activatioa of each of these brain states for each time instance 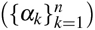 as delineated in (**h**).

Here, we evaluate these two fundamental properties of harmonic brain states, i.e. power and energy, to characterize brain activity in the LSD-induced psychedelic state. We define the power of an harmonic brain state (a connectome harmonic) as the strength its activation for a given time instance in the fMRI data. The energy of a brain state is then estimated as its frequency-weighted contribution to cortical dynamics by combining its power with its intrinsic energy (Methods). Using the introduced connectome-harmonic decomposition, we test whether LSD induced any changes in the energy and power of different frequency connectome harmonics. We further investigate whether the changes in the activation of individual harmonics lead to any variations in the dynamical repertoire of these harmonics brain states. To further characterize the LSD-induced changes in brain dynamics, we evaluate cross-frequency interactions among different connectome harmonics over time. Finally, we also test whether LSD caused any re-organization of brain dynamics at the edge of criticality - i.e. balance between two extreme tendencies; a quiescent order and a chaotic disorder - as hypothesized by previous theoretical studies of this psychedelic state^11^.

## Results

Here we recorded fMRI data from 12 subjects in 6 different conditions; LSD, placebo (PCB), LSD and PCB while listening to music, LSD and PCB after the music session. Exploring the combined effects of music and the psychedelic state induced by LSD provided us an opportunity to reveal not only the LSD-induced dynamical changes in the brain but also how these dynamics are affected by the presence of a complex, natural stimuli like music. Furthermore, music is also known for its capacity to elicit emotions, which is found to be emphasized by the effect of psychedelics^16^. This was a within-subjects design in which healthy participants (mean age 30.5 ± 8, 4 females) received 75 *μ*g LSD in 10 mL saline (intravenous, I.V.) or placebo (10 mL saline, I.V.), 70 minutes prior to fMRI scanning. LSD and placebo sessions were separated by 14 days and occurred in a counter-balanced order, as in^10^.

To study the LSD-induced changes in cortical dynamics, we decomposed fMRI recordings of 12 subjects in 6 different conditions into the activity of frequency-specific brain states (cortical patterns) (Fig. 1**f-h**). The brain states are defined as spatial patterns formed by fully synchronized activity, each associated with a different spatial wavelength; i.e. connectome harmonics^15^ (Fig. 1**f**). We firstly investigated two fundamental properties of these harmonic brain states: 1) power of activation; i.e. the amount of contribution of each harmonic brain state to cortical dynamics, and 2) energy of each of these brain states that combines their intrinsic, frequency-dependent energy with the strength of their activation. Furthermore, to characterize dynamical changes in the repertoire of active brain states, we explored cross-frequency correlations across different harmonics brain states. Finally, to assess the proximity of brain dynamics to criticality, we evaluated power-law distributions across the whole power spectrum of these brain states.

### LSD increases power and energy of brain states

We first estimated the power of activation for each brain state by measuring the strength of its contribution to cortical activity pattern of the fMRI volume acquired at each time instance. By combining the power of activation with the intrinsic, frequency-dependent energy of each harmonic brain state, we calculate the energy of a particular brain state (Methods).

Based on these introduced fundamental measures, enabled by connectome-harmonic decomposition of fMRI data, we first investigated global effects of LSD over the complete spectrum of brain states. To this end, we measured the total power and total energy over the whole connectome-harmonic spectrum for all 6 conditions; LSD, PCB, LSD with-music, PCB with-music, LSD after-music, PCB after-music (Methods). This analysis revealed that the total power and energy of all brain states averaged across all time points significantly 92 increases under LSD (*p* < 0:0001, two-sample t-test between each pair of LSD vs PCB 93 conditions) (Fig. 2**a, b**).

**Figure 2:**
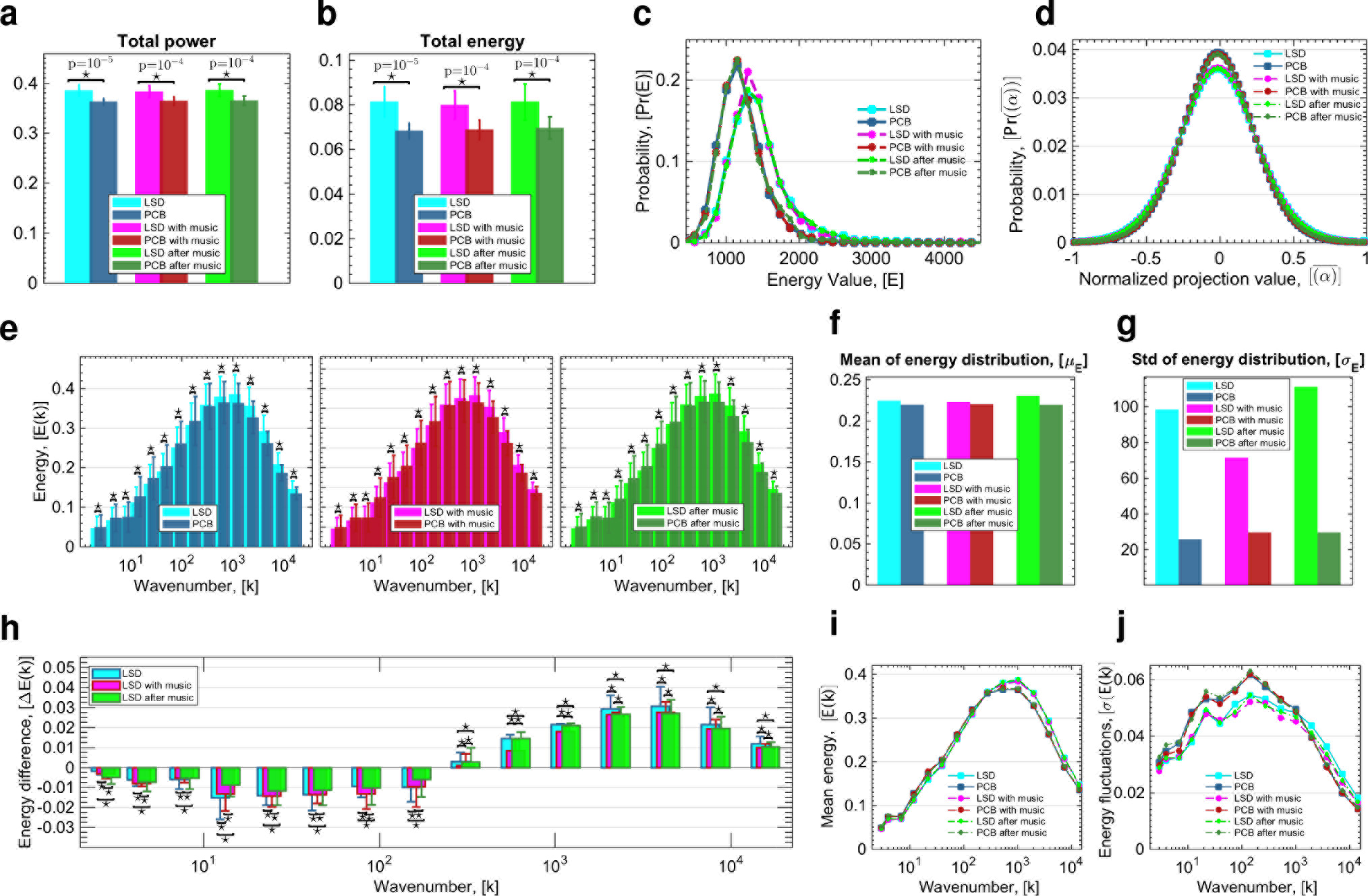
Changes in energy of brain states under LSD. Total power (**a**) and total energy (**b**) of all brain harmonic brain states for all 6 conditions, where stars indicate significant differences (*p*<10^−4^, two-sample t-test) between each pair of LSD vs. PCB conditions with indicated *p*-values. (**c**) Probability distribution of total energy values (sum over all harmonics) for all 6 conditions. (**d**) Probability distribution of the occurrence of projection values (the amount of contribution) of connectome harmonics after normalization of each harmonic’s contribution by the maximum value of the baseline (PCB) condition, shown for all 6 conditions; LSD, PCB, LSD with-music, PCB with-music, LSD after-music, PCB after-music. (*e*) Energy of connectome harmonics quantized into 15 levels of wavenumbers *k* (in the log-scale) for conditions (left) LSD vs. PCB, (middle) LSD with-music vs. PCB with-music, (right) LSD after-music vs. PCB after-music. Stars indicate significant differences (*p* < 0.01, Monte-Carlo simulations after Bonferroni correction). (**f**) and (**g**) show the mean (*μ*) and standard deviation (*σ*) of the fit of the energy distribution of frequencies shown in (**e**) to normal distribution for all conditions, respectively, (**h**) shows the energy differences for each bin between the conditions LSD and PCB, LSD with-music and PCB with-music, LSD after-music and PCB after-music with stars indicating significant differences between conditions no music, with music and after music (*p* < 0.01, Monte-Carlo simulations after Bonferroni correction). Mean (**i**) and standard (deviation (**j**) of energy values of connectome harmonics 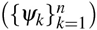 shown as a function of the wavenumber *k*.

To further characterize the energy spectrum of each of the 6 conditions, we estimated the probability distribution of their energy values. We observed significantly different probability distributions of energy values (*p* < 10^−85^, two-sample Kolmogorov-Smirnov test, Fig. S2) between each pair of LSD vs. PCB conditions and reassuringly no significant difference was found between any pair of LSD or any pair of placebo conditions, even in the case where one condition involved listening to music (i.e. only LSD vs. placebo, between-condition differences were found to be significant and not within condition differences). A clear shift to higher energy values was observed at the peak of the probability distribution in all three LSD conditions 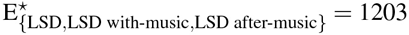 in comparison to the energy probability distributions of PCB conditions 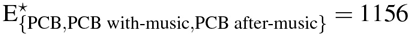 (Fig. 2**c**). In both, the LSD and PCB conditions, with music, we also found a higher probability of reaching this maximum likely (characteristic) energy state 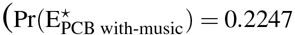 vs. 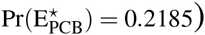 and (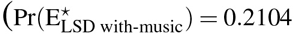 vs. 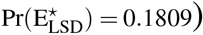. In the placebo condition, this effect of music - increased probability of reaching the characteristic energy state - was also found in the after-music session 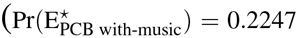 vs. 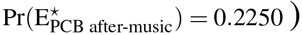. In the LSD after-music condition, the higher probability of reaching the characteristic energy state induced by music, slightly decreased 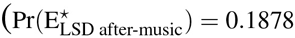 vs. 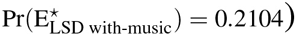, although it still remained higher than in the initial LSD condition (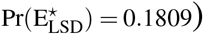, (Fig. 2**c**). These results indicate that LSD renders brain dynamics more likely to reach higher energy states, in particular in response to music, which in turn suggests an increased sensitivity of cortical dynamics to the effect of music. Our results reveal that this amplified effect of music on cortical dynamics was also more rapidly reversed after the offset of music under the influence of LSD compared with the placebo condition. These changes are highly suggestive not only of an increased sensitivity to music, as found in previous studies^16^, but also more rapidly changing cortical dynamics with increased flexibility, which may potentially underlie the enhanced sensitivity to the environment and context observed under the influence of LSD^16–18^. We explore these dynamical changes of cortical activity in further detail in our criticality analysis.

The LSD-induced energy increase of brain activity can be attributed to two possibilities. Firstly, more brain states may be contributing to brain activity leading to an expanded repertoire of brain states, and secondly, the same active brain states may be contributing with more power and energy under LSD. Next, we investigated which of these factors contributed to the observed energy increase under LSD.

### LSD extends the repertoire of active brain states

Theoretical and computational studies indicate that spontaneous brain activity explores a dynamic repertoire of brain states and predict a variation in the size of this repertoire in different states of consciousness^19^. Furthermore, studies exploring psilocybin-induced psychedelic state found greater diversity in functional connectivity motives accompanied by increased variance in temporal oscillations, which indicates an enhanced repertoire of active brain states under the effect of psilocybin, i.e. the main psychoactive compound in magic mushrooms^20^. To quantify the size of the repertoire of active brain states under LSD and placebo conditions, we estimated the probability distribution of the occurrence of projection values (the amount of contribution) of connectome harmonics or all 6 conditions; LSD, PCB, LSD with-music, PCB with-music, LSD after-music, PCB after-music, after normalizing each harmonic’s contribution by the maximum value of the baseline (placebo) condition. Fig. 2**d** demonstrates that the probability distribution of conditions under LSD shows a clear decrease (height of the normal distributions: *μ*_LSD_ = 0.0355, *μ*_PCB_ = 0.0388, *μ*_LSD with-music_ =0.0358, *μ*_PCB with-music_ = 0.0387, *μ*_LSD after-music_ = 0.0356, *μ*_PCB after-music_ = 0.0385) for small magnitude activations (0 value signifying no-activation coincides with the peak of the normal distribution). This decrease signifies that more brain states contribute (with a non-zero weight) to brain dynamics under the influence of LSD. Fig. 2**d** further illustrates the slight increase for higher magnitude activations (towards the tails of the normal distribution, −1 and 1), which indicates that a stronger activation of brain states is observed more frequently under the effects of LSD. The increase in active brain states under LSD is further reflected by the increased width of the normal distribution of projection values (σ_LSD_ = 0.3731, σ_PCB_ = 0.3416, σ_LSD with-music_ = 0.3710, σ_PCB with-music_ = 0.3432, σ_LSD after-music_ = 0.3729, σ_PCB after-music_ = 0.3447) in Fig. 2**d**. These results demonstrate that the increased power and energy of brain activity under LSD is caused by both an extended repertoire of active brain states over time as well increased activity of certain brain states.

### LSD increases high frequency activity

Next we investigated which brain states demonstrate increased activity under the effects of LSD. To this end, we explored frequency-specific alterations in brain dynamics induced by LSD, by first discretising the connectome-harmonic spectrum into 15 levels of wavenumbers *k* in the log-space and then analysing the energy changes within each of these parts of the harmonic spectrum for each of the 6 conditions and each subject separately (Methods).

For all 3 conditions, i.e. before, during and after-music sessions, with LSD vs. PCB, a significant change was observed in the energy of all quantized levels of wavelenths (*p* < 0.01, Monte-Carlo simulations after Bonferroni correction, Fig. 2**e**). Notably, the energy distribution over quantized levels of wavenumbers also followed a log-normal distribution for all conditions (Fig. 2**e**), where both, the mean *μ*[E]), (Fig. 2**f**) and the width (σ[E]), (Fig. 2**g**) of the normal distribution increased in all LSD conditions, although slightly less for LSD with-music condition. Note that the number of divisions of the connectome-harmonic spectrum did not alter the log-normal distribution and the observed energy differences. This increase of the mean and width of the normal distribution suggests that LSD increases the energy (activity) of the brain states corresponding to high frequency wavenumbers. This energy increase for high frequency brain states (connectome harmonics with larger wavenumber *k* > 2 · 10^2^ corresponding to 0.01 − 1% of the whole spectrum) is also clearly observed in Fig. 2**h** demonstrating the energy difference between LSD and PCB conditions before, during and after-music sessions. Critically, a significant decrease of energy is found for all low frequency brain states, connectome harmonics with wavenumer *k* < 2 · 10^2^ corresponding to 0 − 0.01% of the whole spectrum (*p* < 0.01, Monte-Carlo simulations after Bonferroni correction, Fig. 2**h**). For both effects, increased energy of high frequencies and decreased energy of low frequencies, the differences between each pair of conditions were found to be significant (*p* < 0.01, Monte-Carlo simulations after Bonferroni correction). Fig. 2**i** shows the mean energy across all subjects for the discretised spectrum of connectome harmonics and illustrates the increased energy of brain states with larger wavenumbers *k* > 2 · 10^2^. Furthermore, for high frequency brain states (*k* > 10^3^ corresponding to 0.05 − 1% of the whole spectrum) an increase was also found in energy fluctuations over time for all 3 LSD conditions (Fig. 2**j**). Taken together, our results reveal that LSD increases the total energy of brain activity and expands the repertoire of active brain states by significantly increasing the activity of high frequency brain states.

### Cross-frequency correlations between brain states

We next sought to understand whether LSD-induced expansion of the repertoire of active brain states occurred in a structured or random fashion. To this end, we investigated LSD-induced changes in cross-frequency interactions in brain dynamics. We examined the degree of co-activation of different frequency brain states by exploiting the spectra-temporal representation enabled by the connectome-harmonic decomposition. As this harmonic decomposition of fMRI data yields the strength of activation of different frequency brain states over time, the correlation between the time courses of different connectome harmonics reveals the degree to which these two frequency brain states co-activate within the complex cortical dynamics. In this manner, we estimated cross-frequency correlations between each pair of brain states across all LSD and placebo conditions. Under the influence of LSD for all three conditions; before music, with music and after music, we observed a significant decrease in cross-frequency correlations within the low-frequency brain states (*k* ∈ [0 − 0.01%] of connectome-harmonic spectrum) (effect size; Cohen’s *d*-value ⋆ > 0.2, Fig. 3**a**). Notably, this range of brain states is the same as that in which a significant decrease in energy was observed (Fig. 2**h**). In a higher frequency range *k* ∈ [0.01 − 0.1%], the only significant difference was found between LSD and PCB conditions with music (effect size; Cohen’s *d*-value ⋆ > 0.2, Fig. 3**b**) indicating the influence of music on the co-activation of brain states within this frequency range. For increasing frequency, *k* ∈ [0.1 − 0.2%] of the spectrum, no significant differences were found in cross-frequency correlations (Fig. 3**c**). Finally, for higher frequency range *k* ∈ [0.2 − 1%], we found a significant increase in the cross-frequency correlations between LSD and PCB conditions, while this effect was not significant for conditions with and after-music (Fig. 3**d**). Also, over the complete spectrum of brain states, LSD significantly increased cross-frequency correlations (Fig. 3**e**).

Considering the sequential acquisition of the scans in conditions before music, with music and after music, the insignificant differences of the cross-frequency correlations within the high frequency range between LSD and PCB with music and after music can be attributable to both, the effect of music and the fading effect of LSD over time. To distinguish the effect of these two factors, we compared the average cross frequency correlations across the whole connectome-harmonic spectrum between each pair of the 6 conditions (Fig. 3**f-n**). Fig. 3**f** clearly demonstrates the decreased cross-frequency correlations among the low frequency brain states (*k* ∈ [0 − 0.1%] of the spectrum) accompanied by increased correlation between all frequencies in the range *k* ∈ [0.1 − 1%] of the spectrum. Over time, the LSD-induced increase in cross-frequency co-activation gradually diminished (Fig. 3**f-h**), which was also confirmed by the comparison of the sequentially acquired LSD scans (Fig. 3**i-k**). These changes were not found in the sequential comparison of PCB scans (Fig. 3**l-n**). Music under the influence of LSD decreased the cross frequency correlations also within the frequency range *k* ∈ [0.01 − 0.02%] of the spectrum (Fig. 3**g**), which remarkably coincides with the range of brain states whose energy changes were altered under the influence of music (Fig. 2**h**). Notably, this effect of music was observed only in the LSD but not the PCB condition (Fig. 3**m**). This analysis confirms that both factors, the fading effect of LSD and the influence of music, contribute to observed changes in cross-frequency correlations over the three scans (LSD/PCB, LSD with-music/PCB with-music and LSD after-music/PCB after-music). However, while music affected the communication within the low to mid range frequencies *k* ∈ [0.01 − 0.02%] in particular under the influence of LSD, the isolated effect of LSD was apparent in the increased cross-frequency correlations throughout the connectome-harmonic spectrum.

These results demonstrate that for the low-frequency range, where the energy of brain states decrease under the influence of LSD, there is also a decrease in the “communication” (co-activation) of these brain states. The exact opposite effect, i.e. increased communication along with increased energy and power, is observed among a large part (*k* ∈ [0.01 − 1%]) of the spectrum under the influence of LSD. Such increased cross-frequency correlations strongly suggest that LSD causes a re-organization rather than a total randomization of brain dynamics. This type of non-random expansion of the state repertoire naturally occurs in dynamical systems when they approach criticality - the boundary of an order-disorder phase transition^21,22^. As a logical next step, we therefore investigated whether the dynamics of harmonic brain states show other characteristics of criticality and how these may be altered under the effect of LSD.

### Power laws and whole-brain criticality

With a growing body of experimental evidence^23–30^and theoretical findings^31–37^, it has become increasingly apparent that neural activity shows characteristics of criticality - a delicate balance between two extreme tendencies; a quiescent order and a chaotic disorder - where complexity increases and certain functional advantages may emerge^38,39^. Theoretical and computational studies identify that criticality enables the essential dualism necessary for complex dynamics; i.e. a certain level of stability (order) is required for coherent functioning and certain degree of disorder is needed for functional flexibility and adaptability^34^. These studies also highlight some important functional advantages of criticality; e.g. that greater diversity in the repertoire of brain states^22^ enables a larger capacity for information encoding^22^ and faster information processing^22,30^.

Supporting the hypothesis that brain dynamics reside at the edge of criticality, experimental studies reveal a key characteristic of critical dynamics - the power-law distributions - in large scale brain networks in fMRI^24,27^, electroencephalography (EEG)^23,26,28^, MEIG^23,24,28,29^and intracranial depth recordings in humans^40^ as well as in numerical simulations of computational models of brain dynamics^33,35^, mostly with small deviations from criticality to the subcritical (ordered) regime.

The power-laws, although observed consistently across wakefulness^40^, deep-sleep^40^, REM sleep^40^ and during anaesthetics induced loss of consciousness^25^, are found to slightly deteriorate in wakefulness^30,37,40^, tend to diminish in cognitive load^41^ and recover during sleep^32,37,40,42^. These findings suggest that even though power-laws are likely to be a feature of neural dynamics, which transcends levels of consciousness, differences in power-law distributions are characteristic of different states and the proximity of these states to critical dynamics. Furthermore, such deviations and subsequent re-emergence of power-laws with changing states of consciousness and cognitive-load strongly indicate that they originate from critical network dynamics, ruling out alternative explanations such as filtering or noise^37^.

In line with these findings, the tuning of brain dynamics towards or away from criticality is likely to be mediated by varying the excitation/inhibition balance^37,43,44^, which has been shown to underlie the temporal organization of neuronal avalanches with power-law distributions^43^ as well as whole-brain oscillatory patterns^15^. In contrast, other pharmacologically induced variations in neural activity; e.g. changes in the concentration of dopamine or administration of a dopamine D1 receptor agonist,^45^ or antagonist^46^, as well as application of acetylcholine^47^, lead to alterations of the steepness of the critical exponent without destroying the power-law distributions. As with other classic psychedelic drugs^48^, LSD’s principal psychoactive effects are mediated by serotonin 2A receptor agonism, and 5-HT2AR signalling has reliably been shown to induce cortical excitation^49^. Increased cortical excitation via increased 5-HT2AR signalling is a plausible mechanism by which LSD may tune cortical dynamics towards criticality.

**Figure 3.**
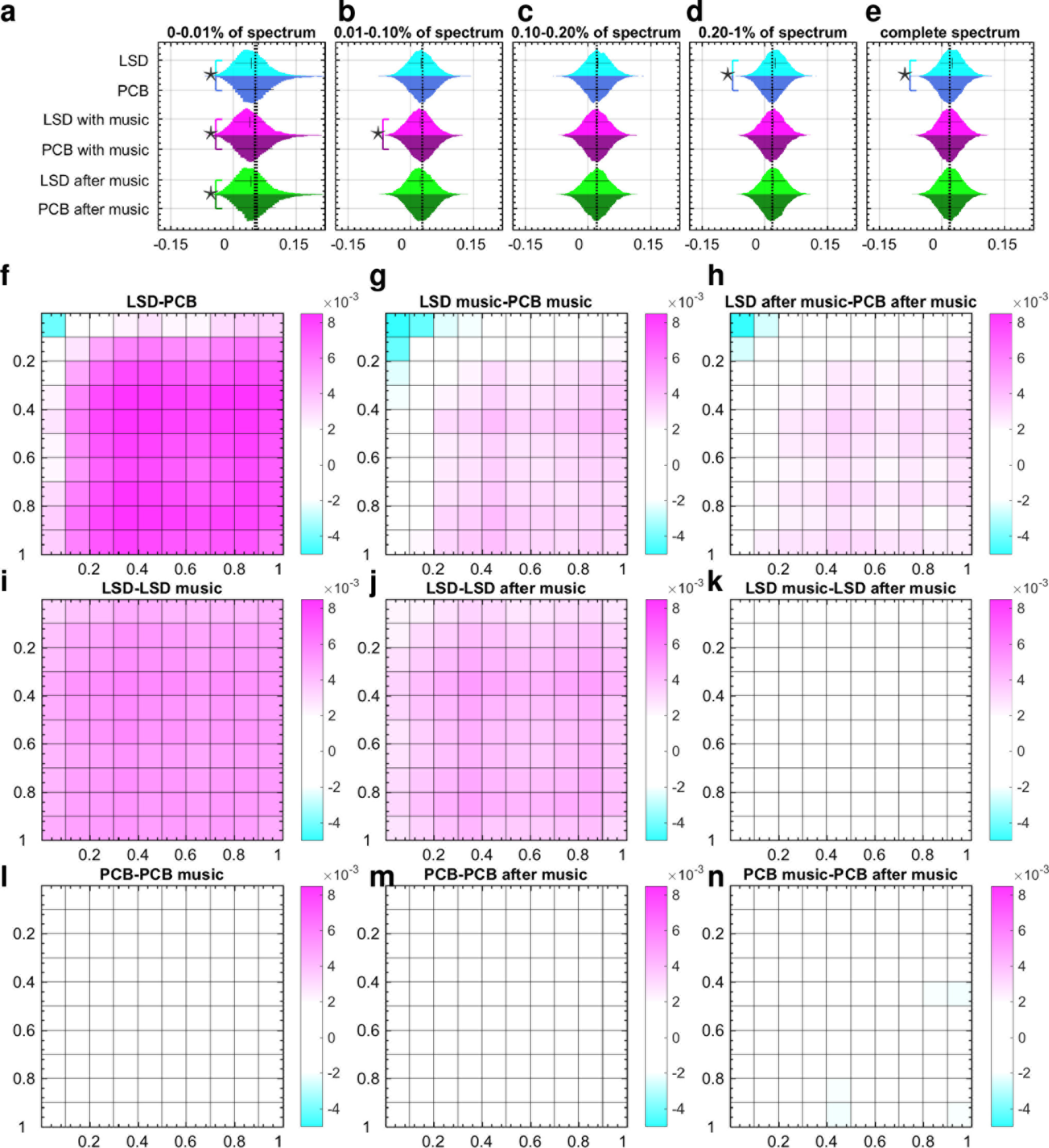
Cross frequency correlations. (**a**)-(**d**) Distributions of cross-frequency correlation values within [0 − 0.01%], [0.01 − 0.1%], [0.1 − 0.2%] and [0.2−1%] of the spectrum, respectively, (**e**) shows the distribution of cross-frequency correlations across the complete spectrum of connectome harmonics. Significant differences between cross-frequency correlation distributions are marked with stars (effect size; Cohen’s *d*-value ⋆ > 0.2) for pairs of condition LSD, PCB, LSD with-music, PCB with-music, LSD after-music, PCB after-music. (**f**)-(**n**) illustrate differences in mean cross frequency correlations in 10 × 10 partitions across the complete spectrum of connectome harmonics evaluated between all pairs of 6 condition; LSD, PCB, LSD with-music, PCB with-music; LSD after-music, PCB after-music.

Based on LSD’s known pharmacology and related effects on cortical excitation^10^, as well as a prior hypothesis regarding the psychedelic state and criticality in the brain^11^, we investigated LSD induced dynamical changes in the brain in the context of criticality. To this end, we evaluated the distribution of maximum power, average power as well as power fluctuations over the spectrum of connectome harmonics. Notably, all power related distributions (maximum, mean and standard deviation) of the connectome harmonics with different wavenumbers followed power-law distributions (Fig. 4). In line with previous findings^30,32,40,42^ and theoretical models^31–37^, we observed a slight cut-off in the tail of the power-law distributions indicative of the slight deviation to the subcritical regime. This result confirms previous studies suggesting that conscious, waking state brain dynamics reside at the edge of criticality with a slight deterioration to the subcritical regime^35,37,40^.

In an effort to quantify LSD-induced changes to critical brain dynamics, we quantitatively evaluated the goodness of fit of power-laws, by measuring the root mean squared error ε of power-law fit for all different conditions (Methods). Critically, the root mean squared error of power-law-fits decreased significantly (*p* < 0.01, two-sample t-test) for maximum power (Fig. 4**a-b**), mean power (Fig. 4**d-e**) and power fluctuations (Fig. 4**g-h**) in LSD compared with PCB for the first two conditions (before music and with music). The decreased error of fit found for LSD vs. PCB in the after-music condition (Fig. 4**c,f,i**) remained insignificant, reflecting the slightly fading effect of LSD over the course of the three scans. The decreased goodness-of-fit error demonstrates that the distribution of all power-related observables (maximum power, mean power and power fluctuations) fit power-law distribution more closely under the influence of LSD. These experiments suggest that brain dynamics in both conditions, LSD and PCB, reside close to criticality with slight deviations to the subcritical regime, as also indicated in previous studies^30,37,40^, while the induction of LSD tunes brain dynamics further towards criticality.

An additional analysis evaluated the power-law exponent for all 6 conditions. In all conditions with LSD compared to placebo, the power law exponent of maximum (Fig. 4**a-c**) and mean-power distribution (Fig. 4**d-f**) is found to decrease significantly (*p* < 0.01, two-sample t-test). For power fluctuations, the decrease was only significant for the first scan (Fig. 4**g**); LSD vs. placebo condition, coinciding with the peak of the LSD experience. This change in the power-law exponents under the influence of LSD indicates increased power and power-fluctuations of high frequency and slightly decreased power and power-fluctuations of low frequency connectome harmonics. As the decrease in power-law exponent and goodness-of-fit error both originate from the increased power of high frequency connectome harmonics, this finding confirms our earlier results regarding increased energy in high frequency states and enriched repertoire of brain states under the influence of LSD while indicating a crucial link between whole-brain criticality and the observed energy, power and repertoire changes.

### LSD-induced energy changes correlate with subjective ratings

We also investigated how the LSD-induced changes in brain activity relate to subjective experience. Participants were asked to perform a limited number of visual analogue scale (VAS) style ratings at the end of each scan, using a button box in the scanner. Five key facets of the LSD experience were enquired about: 1) complex imagery (i.e. eyes-closed visions of objects, entities, landscapes etc.), 2) simple hallucinations (i.e. eyes-closed visions of shapes, colours, geometric patterns etc.), 3) emotional arousal (i.e. how emotional the participant felt, regardless of whether emotions were positive or negative), 4) positive mood, and 5) ego-dissolution (i.e. a fading sense of self, ego and/or subjectivity).

To examine the relation between the activation of different brain states and subjective experiences, we explored the correlations between energy changes of different frequency connectome harmonics and subjective ratings of the five experiences. We first estimated the amount of change in energy between LSD and PCB conditions for different connectome harmonics across all 12 subjects for the two scans without music (LSD/PCB and LSD after-music/PCB after-music). Then, we evaluated the correlations between the subjective ratings and the estimated energy differences ΔE = (E_PCB_ − E_LSD_) for both, individual harmonics and different ranges of the harmonic spectrum.

For the low frequency range *k* ∈ [1 − 200] corresponding to [0 − 0.01%] of connectome-harmonic spectrum, we observed generally a *decrease* in the mean energy as well as in the energy fluctuations under LSD at an individual subject level, as indicated by the positive values of 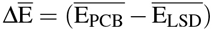 and Δσ (E = (σ (E_PCB_) − σ(E_LSD_)) denoting the differences in the mean and in the standard deviation of energy between the placebo and LSD conditions, respectively (in Fig. 5**a, b, d, e**). The amount of the decrease in the mean energy 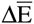 of this frequency range significantly correlated with the subjective ratings of ego-dissolution (*r* = 0.55317, *p* < 10^−3^, Fig. 5**a**) and emotional arousal (*r* = 0.61063, *p* < 10^−3^, Fig. 5**b**). We found similar correlations also between these subjective experiences and energy fluctuations of the same range of connectome harmonics (Fig 5**d, f**). These findings suggest that the deactivation of the low-frequency connectome harmonics play a crucial role in the neural correlates of ego-dissolution and emotional arousal. Remarkably, this part of the connectome harmonic spectrum also corresponds to the same frequency range in which decreased energy (Fig. 2**h**) and decreased cross-frequency correlations (Fig. 4) were found under LSD.

**Figure 4.**
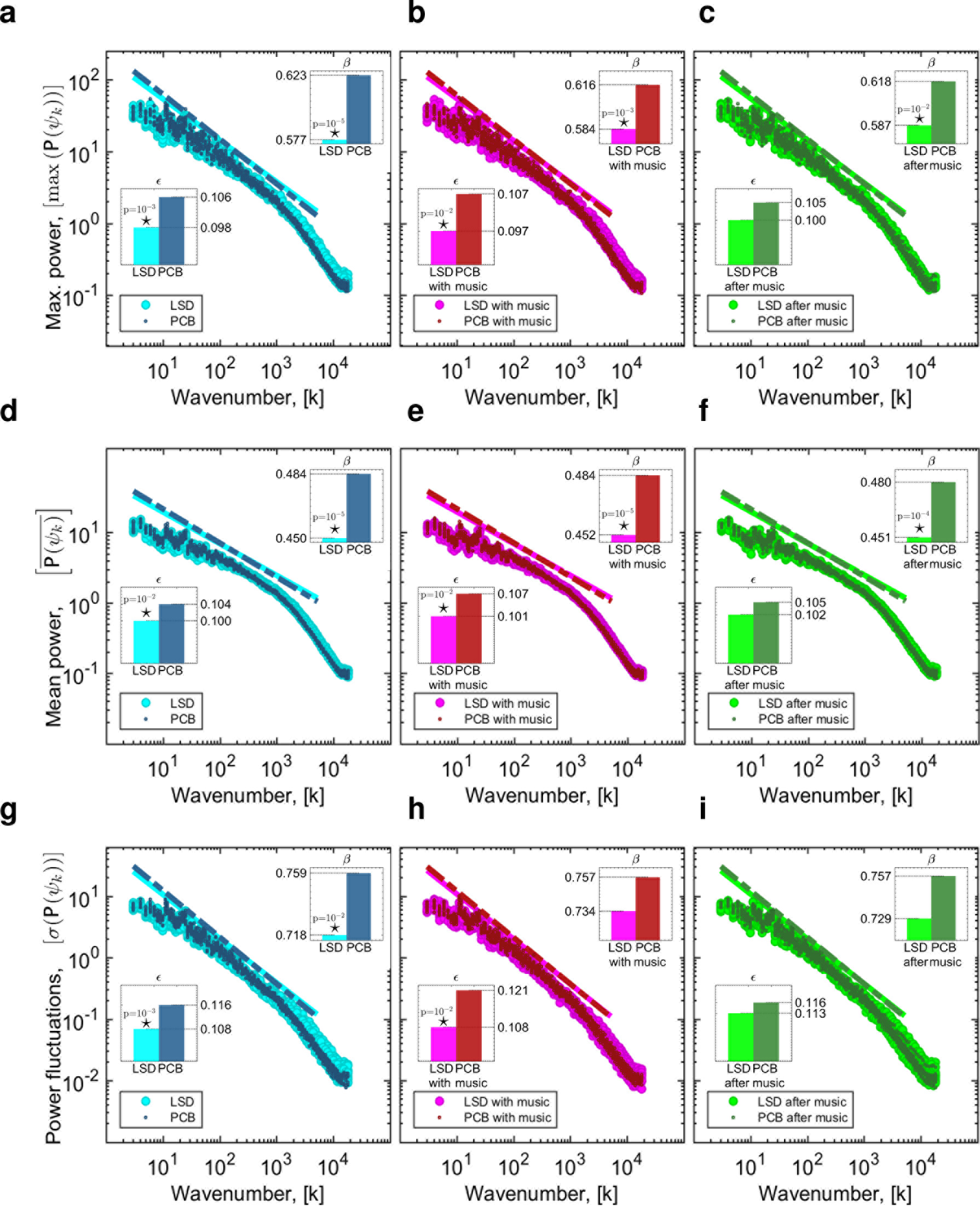
Power laws in connectome harmonic decomposition. Maximum power (max(P(*ψ_i_*))) vs. wavenumber (*k*) of connectome harmonics 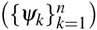 in log_10_ coordinates for (**a**) LSD vs. PCB, (**b**) LSD with-music vs. PCB with-music and (**c**) LSD after-muric vs. PCB after-music, respectively. Mean power 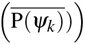 vs. wavenumber (*k*) of connectome harrmonics 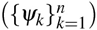 in log_10_ coordinates for (**d**) LSD vs. PCB, (**e**) LSD with-music vs. PCB with-music and (**f**) LSD after-music vs. PCB after-music, respectively. Power fluctuations (σ(P(*ψ*_*k*_))) vs. wavenumber (*k*) of connectome harmonics 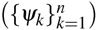 in log_10_ coordinates for all 6 conditions. In all plots, *ε* and *β* indicate the root mean squared error and the slope of the line fit, respectively. Stars indicate significant differences (*p* < 0.05, two-sample t-test).

The energy change within this low frequency range alone did not significantly correlate with ratings of positive mood under LSD. But a broader range of spatial frequencies *k* = [1, ···, 1100] showed significant correlation with the intensity of positive mood induced by LSD (*r* = 0.45629, *p* < 10^−2^ for 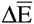 and *r* = 0.4714, *p* < 10^−2^ for Δσ(E)). This finding implies that only a decrease of activity of low frequency connectome harmonics *k* = [1, ···, 200] is not sufficient to account for the positive mood felt under LSD, but this decrease has to be accompanied by increased contribution of slightly higher frequency range connectome harmonics *k* = [201, ···, 1100] for positive mood to be felt. Note that LSD generally induced a decrease in the low frequency range *k* = [1, ···, 200] while leading to increased activation in the rest of the connectome harmonic spectrum *k* = [200, ···, 18715], as shown in Fig. 2**h**.

For individual harmonics, although we observe high correlations ranging from −0.6 to 0.5 between the energy change of each harmonic and the subjective ratings, these correlations did not survive a conservative correction for multiple comparisons (Bonferroni correction), when the whole connectome-harmonic spectrum was considered (18715 comparisons, Fig. S3 shows the correlation values of the first 200 harmonics). We did not find any significant correlations between the subjective ratings of simple hallucinations and complex imagery and the energy changes of a particular frequency range of brain states.

**Figure 5.**
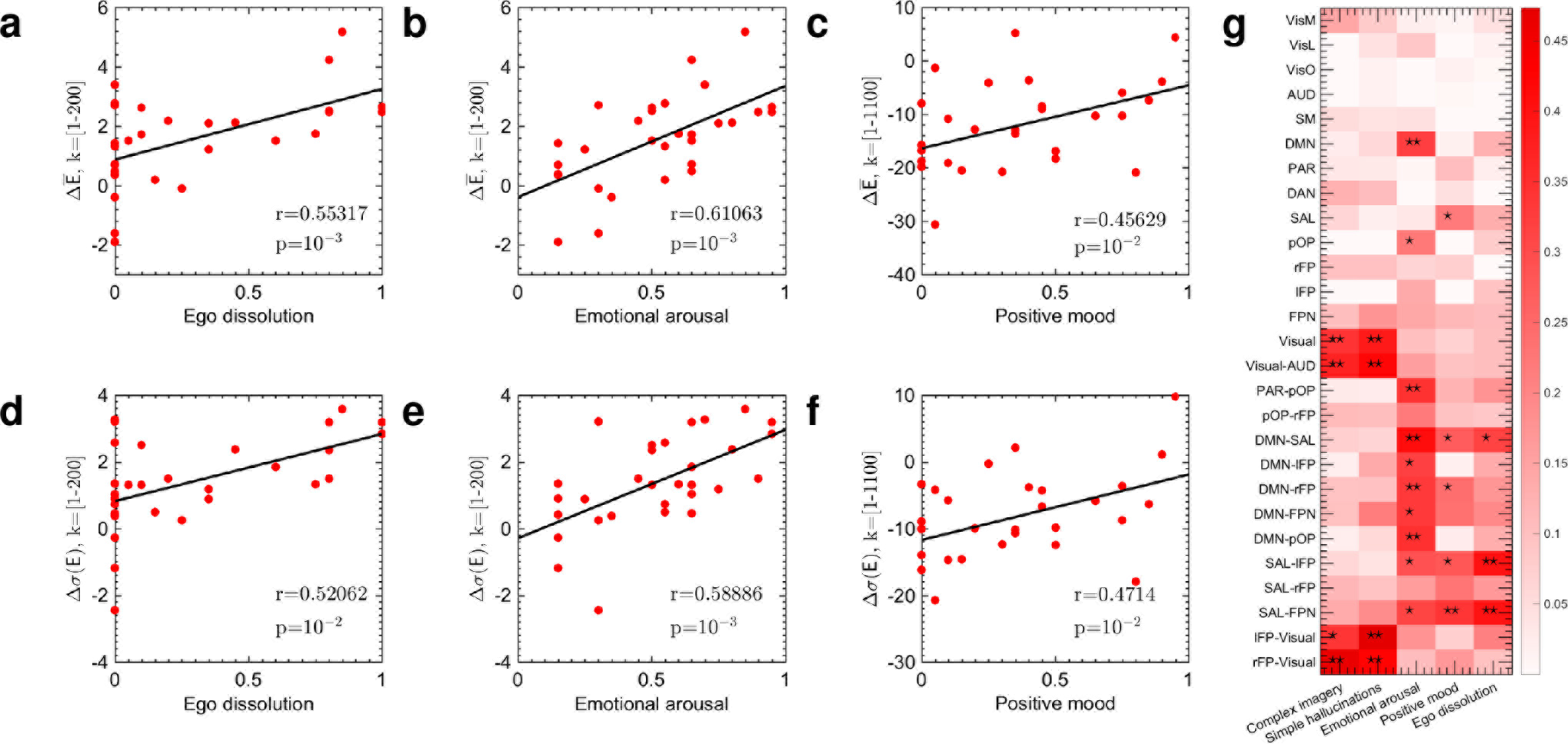
Correlations between energy changes of connectome harmonics and subjective experiences. (**a**) and (**b**) demonstrate significant correlations between the difference in mean energy of connectome harmonics 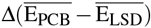 for low frequency connectome harmonics *k* = [1, ···,200] and the subjective ratings of ego dissolution and emotional arousal, respectively. (**c**) shows the correlation between the energy difference of connectome harmonics 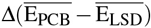 for a broader frequency range of connectome harraonics *k* = (*r*, ···, 1100] and the subjective ratings or positive mood. (**d**) and (**e**) demonstrate significant correlations between the difference in energy fluctuations of connectome hcrmonics Δ(σ(E_PCB_) − σ(E_LSD_)) for low frequency connectome harmonics *k* = [1, ···,200] and the subjective ratings of ego dissolution and emotional arousal, respectively, (**f**) shews the correlation between difference in energy fluctuations of connectome harmonics Δ(σ(E_PCB_ − σ((E_LSD_)) for *k* = [1, ···, 1100] and the subjective ratings of positive mood. (**g**) illustrates multiple correlations between the functional connectivity changes of groups of resting state networks (RSNs) and subjective experiences estimated using 200 brain states, *k* = [1, ···,200] *p*: ⋆ < 10^−10^,*p*: ⋆⋆ < 10^−15^ after Bonferroni correction. Correlation strengths are represented by the intensity of red for each pair.

### Correlations between subjective ratings and connectivity of resting state networks

Finally, we asked whether there was a correlation between subjective ratings and connectivity of testing state networks (RSNs). Previous analyses with LSD^10^ and other psychedelics^11,50,51^ have shown changes in the functional properties of RSNs under these drugs that correlate with different psychological aspects of the experience.

Even for simple hallucinations, it is now understood that no one single brain area is responsible, but rather the interaction of multiple brain areas^52^. Thus, unlike previous studies focusing on the connectivity of individual networks^10,11,50,51^, here we investigated how the connectivity changes in groups of networks mutually relate to the intensity of subjective experience using multiple correlation analysis^53^. We examined the correlations between the ratings of subjective experiences of simple hallucinations, complex imagery, emotional arousal, positive mood and ego dissolution and LSD-induced functional connectivity changes of RSNs, individually, as well as in subgroups of RSNs using multiple correlation analysis^53^.

For the group analysis we first identified the following RSNs as described in^10^: medial visual network (VisM), lateral visual network (VisL), occipital pole network (VisO), auditory network (AUD), sensorimotor network (SM), default-mode network (DMN), parietal cortex network (PAR), dorsal attention network (DAN), salience network (SAL), posterior opercular network (POP), left fronto-parietal network (lFP) and right fronto-parietal network (rFP) and then defined the following subgroups of networks: fronto-parietal network (FPN by combining lFP and lFP), visual (by combining VisM, VisL, VisO), visual-AUD, PAR-pOP, pOP-rFP, DMN-SAL, DMN-lFP, DMN-rFP, DMN-FPN, DMN-pOP, SAL-lFP, SAL-rFP, lFP-visual, rFP-visual.

As connectome harmonics are spatial patterns of synchronous activity emerging on the cortex for different frequency oscillations, they are theoretically equivalent to frequency-specific functional connectivity patterns^15^; hence, the observed functional connectivity changes in the RSNs can be attributed to and decomposed into the changes in the activation of individual connectome harmonics. Here, we evaluated multiple correlations between groups of RSNs and subjective ratings using both, estimated functional connectivity changes of RSNs directly and their correlations with the energy changes of individual harmonics (Methods).

Although the evaluation of the multiple correlation analysis between subjective ratings and connectivity changes of RSNs showed differences of correlations (Fig. S4a), in this direct application, no correlation was found to be significant. One limitation of this approach is that the small number of data points (12 subjects × 2 scans leading to 24 dimensional vectors to compute the multiple correlations) renders the correlation analysis poorly powered to reveal potentially ‘true’ relations between RSNs and subjective ratings (i.e. the risk of false negatives). This limitation can be addressed by firstly correlating the connectivity changes of RSNs to energy changes of individual brain states (connectome harmonics) and then evaluating the multiple correlations between the subjective ratings and individual or groups of RSNs (Methods). This approach has the advantage of enabling hidden information present in the data itself to emerge by revealing the contribution of each brain state to changes in functional connectivity. The number of brain states chosen to express the connectivity changes of RSNs will determine the sensitivity of multiple correlations (Fig. S4, Methods). Fig. S4 shows the multiple correlation values evaluated on the RSN connectivity directly, and for increasing number of brain states (30, 50, 100, 200 and 18715 (complete spectrum)). Although, similar correlation matrices emerge in the direct (on the RSN connectivity) and indirect (on the energy changes of connectome harmonics) evaluation of multiple correlations, the latter revealed significant correlations between the intensity of certain subjective experiences and groups of RSNs.

Firstly, we observed significant correlations between the connectivity changes of visual and sensory (visual-auditory) networks and ratings of simple hallucinations and complex imagery (Fig. 5**g**, ⋆⋆: *p* < 10^−15^, *k* ∈ [1, ···, 200], after Bonferroni correction). However, this significant correlation was only observed when the visual or sensory networks are considered together but not individually. This finding suggests that it is not the activity of individual networks alone but rather their joint activity that relates to the experience of hallucinations and imagery. Moreover, the coupled connectivity of the the visual networks with the left fronto-parietal network (lFP) correlated with ratings of simple hallucinations (Fig. 5**g**, ⋆⋆: *p* < 10^−15^, *k* ∈ [1, ···, 200], after Bonferroni correction). Interestingly, the coupled connectivity of the right fronto-parietal network (rFP) with the visual networks was found to significantly correlate with the ratings of both, complex imagery and simple hallucinations (Fig. 5**g**, ⋆⋆: *p* < 10^−15^, *k* ∈ [1, ···, 200], after Bonferroni correction) suggesting that the asymmetric contribution of the fronto-parietal networks may underlie the perceptual abnormalities such as visual hallucinations experienced in the psychedelic state.

Secondly, we also found significant correlations between the DMN connectivity and the intensity of emotional arousal (Fig. 5**g**, ⋆⋆: *p* < 10^15^, *k* ∈ [1, ···, 200], after Bonferroni correction). The changes in coupled connectivity of DMN with the salience network (SAL) - a network of brain areas that plays an important role in attentional capture of biologically and cognitively relevant events^52^ - showed significant correlations with all three experiences of emotional arousal, positive mood and ego dissolution (Fig. 5**g**, ⋆⋆: *p* < 10^−15^, *k* ∈ [1, ···, 200], after Bonferroni correction). Importantly, abnormal DMN-SAL functional connectivity correlating with intense subjective effects has previously been reported under LSD^10^ and psilocybin^54^. We also found that the coupled connectivity changes of SAL with lFP alone or FPN (lFP and rFP together) significantly correlated with the ratings of emotional arousal, positive mood and ego-dissolution, whereas connectivity changes of the SAL when coupled only with rPF did not yield the same level of significance (Fig. 5**g**, ⋆: *p* < 10^−10^, *k* ∈ [1, ···, 200], after Bonferroni correction) for correlations with the ratings of these experiences. In particular, the coupling of the the rFP and lFP together with the SAL increased the significance of the correlations with positive mood (Fig. 5**g**, ⋆⋆: *p* < 10^−15^, *k* ∈ [1,···,200], after Bonferroni correction). The coupled connectivity changes of posterior opercular network (pOP) and DMN as well as pOP and the parietal network (PAR) also showed significant correlation with the ratings of emotional arousal, whereas the the correlation of pOP connectivity alone was less significant (Fig. 5**g**, ⋆: *p* < 10^−10^, *k* ∈ [1, ···, 200], after Bonferroni correction). All four pairs of networks, DMN-SAL, DMN-rFP, DMN-pOP, PAR-pOP, which significantly correlated with the ratings of emotional arousal, have been previously reported to show increased functional connectivity under LSD^17^ and the DMN specifically has been linked to mood and emotion^52,55–58^as well as ego-dissolution in relation to psychedelics^10,11^. Our findings suggest a potential link between the increased between-RSN functional coupling, particularly in relation to high-level RSNs, and emotional arousal under the effect of LSD. Further important parallels emerge between these findings linking the RSNs to the intensity of subjective experiences in the psychedelic state and the abnormal activity of RSNs in psychiatric disorders, as explained in Discussion.

## Discussion

Here, we investigated LSD-induced changes in brain activity using a novel connectome-specific harmonic decomposition method. In particular, utilizing the connectome harmonics^15^ as brain states - elementary building blocks of complex cortical activity - we studied LSD-induced changes in energy and power of these harmonic states as well as the dynamical changes in their active repertoire.

Unlike previous techniques applied to explore brain dynamics such as dynamic functional connectivity^59^, the estimation of harmonic brain states relies solely on the structural connectivity (human connectome) and thus is independent from the fMRI data itself. Hence, the connectome harmonics are estimated once as the harmonic modes of the human connectome and serve as a universal, anatomically informed harmonic representation to explore any functional neuroimaging data; i.e. similar to using sine and cosine functions to decompose a signal in Fourier transform. This approach allows us to use the exact same brain states to describe the brain dynamics in different conditions; i.e. LSD vs. placebo, which would not be possible with previous techniques. Furthermore, by definition, the connectome harmonics provide fully synchronous (spatial) patterns of activation, each spatial pattern corresponding purely to a different temporal frequency oscillation^15,60^. This allows for a frequency-specific decomposition of the fMRI data for the first time in the spatial domain. Finally, although in this study we apply this technqiue to fMRI data in order to extract the neural signatures of the LSD state, application of the connectome-harmonic decomposition can also be easily extended to other functional neuroimaging datasets such as MEG or high density scalp EEG.

Our results demonstrate that LSD alters the energy and the power of individual harmonic brain states in a frequency-selective manner and enriches the connectome-harmonic repertoire. Moreover, this expansion of the repertoire of active brain states occurs in a non-random fashion with increased co-activation across frequencies suggesting a re-organization of brain dynamics. Taken together, the expanded repertoire of brain states and increase in cross-frequency correlations under LSD demonstrate that not only do *more* brain states contribute to neural activity under LSD leading to a richer, more flexible repertoire of dynamics, but also their co-activation patterns are highly correlated over time, indicating a preserved stability in brain dynamics, albeit a ‘stability’ of a different kind with more complex dynamics.

Increased diversity in the repertoire of brain states is an expected property of brain dynamics approaching criticality - i. e. balance between order and chaos^22^. Such an expansion of brain states is thought to underlie an expanded capacity for information encoding^22^ and an enhanced efficiency of processing^22,30^. In light of previous theoretical^11^ and experimental findings^21,23–30^, we hypothesized that this increased energy and enriched repertoire of brain states under LSD could be accompanied by a tuning towards criticality. Confirming this hypothesis, we found that LSD induced both: closer fit of power-laws in the distribution and fluctuations of several observables and a slight change in the critical exponent, indicating a shift of brain dynamics towards criticality. Our results suggest that brain dynamics in both conditions, LSD and placebo, reside close to criticality, with slight deviations to the subcritical regime under placebo, as also indicated for resting state brain dynamics in previous studies^30,37,40^, while the induction of LSD tunes brain dynamics further towards criticality. It is noteworthy that the brain dynamics at rest have been found to fluctuate near criticality rather than sitting at a singular critical point^27,33^. These theoretical^33^ and empirical findings^27^ imply the presence of an extended critical region, which was shown to precisely stem from the hierarchical organization of cortical networks^39^. Our results reveal that while the hallmark of critical dynamics, the power-laws, were observed in both the LSD and placebo conditions, the significantly reduced goodness-of-fit error of power-laws under LSD, suggests a tuning of brain dynamics further towards criticality - consistent with the so-called ‘entropic brain’ hypothesis^11^.

The presented method goes beyond the conventional fMRI analyses previously used to measure changes in brain activity under psychedelics^10,61^, by enabling the study of fundamental properties of harmonic brain states, such as energy and power, and by revealing how these introduced concepts relate to important principles of dynamical systems, such as whole-brain criticality. Criticality in the brain (i.e. the property of being optimally poised between order and disorder) has been hypothesised to reflect and underlie its advanced functionality^33,38^; remarkably however, here, using the above-described connectome-harmonic decomposition, we discover a tonic brain state (the LSD state), which exhibits more pronounced signatures of criticality than the normal waking state. This finding provides the first experimental evidence for previous theoretical studies hypothesizing that brain activity in the psychedelic state may be closer to critical dynamics than the normal waking state^11^ and has several important implications:

Firstly, criticality provides the necessary conditions for optimal information processing and representation^22^ for a complex system, rendering it more supple and flexible within its own intrinsic functioning but also more sensitive to incoming stimuli^22,37^. Hence, a natural functional consequence of tuning brain dynamics towards criticality, as was observed here under LSD, is an increased sensitivity to both external stimuli as well as internal, intrinsic activity, which in turn leads to greater sensitivity to both the external environment and internal milieu - referred to as ‘setting’ and ‘set’ respectively, in relation to psychedelics^18^. Hence, the LSD-induced shift towards criticality, presents a candidate mechanism underlying increased sensitivity to the context under LSD and psychedelics more generally^17^. The importance of so-called ‘set and setting’ have been much emphasized by those working with psychedelics in humans^18,62^and the very definition of the word ‘psychedelic’ is intended to refer to the putative ability of these compounds to allow the surfacing of normally ‘unconscious’ mental contents into consciousness^63^. The present findings may therefore represent the beginnings of a mechanistic explanation of these principles, which if substantiated, would have profound implications for psychology.

Secondly, various previous studies have pointed out that deviations from criticality could be symptomatic or even causative of certain psychiatric disorders^22,36,37,64^. In particular, brain dynamics in depression^11,32^, addiction^11^ and obsessive compulsive disorder (OCD)^11^ have been associated with the subcritical regime^11,32^, whereas super-critical regime has been found to govern brain dynamics during epileptic seizures^30,32,37,64^ and in conditions such as autism^22^. Taken together with these studies, our findings highlight the potential effect of LSD to regulate brain dynamics in pathology by re-establishing the critical balance between ordered and disordered states. Such an action may explain the increasing body of evidence supporting the therapeutic potential of psychedelic drugs for treating disorders such as OCD^65^, depression^66,67^ and addictions^68^.

Finally, the balance between the complementary dynamics governing stability (ordered regime) and flexibility (disordered regime), attained at criticality, enables flexible and evolving dynamics while maintaining stability. Thus, brain dynamics at the edge of criticality have been hypothesized to constitute the neural basis of creativity^69^. Supporting evidence of this hypothesis comes from studies revealing network correlates of creativity^70^. Divergent thinking and creative idea production have been found to involve the cooperation of two different types of brain networks: those linked to top-down control of attention and cognition (SAL, executive control network (ECN)) and the DMN associated with spontaneous thought^70^. This finding resonates with the above-mentioned functional advantages of a critical system, where an optimal balance between stability (cognitive control) and flexibility (spontaneous thought) may enable the generation of novel and potentially useful ideas.

An intuitive understanding of the relation between creativity, critical dynamics and the connectome-harmonic decomposition method utilized here, can be gained from studies exploring neural basis of jazz improvisation^71^. A notable finding of these studies is that the number of musical notes played during improvisation is significantly higher compared with memorized play of the same piece, hence leading to an increase of novel information^71^; i.e. improvisation (involving creativity) introduces spontaneously generated novelty into previously known patterns of melody. Likewise, brain dynamics at the edge of criticality enable the emergence of maximally novel dynamics, where more harmonic brain states are involved in a structured (non-random) yet spontaneous manner, as demonstrated in our findings. Note that connectome harmonics, utilized as brain states in this work, are also mathematically equivalent to the patterns of standing sound waves emerging within musical instruments, where in this case the standing wave equation is solved for the particular connectivity of the human brain (connectome)^15^. Consistent with this hypothesis, our findings reveal both, an increase in the number of active brain states, accompanied by a shift towards criticality in brain dynamics under the effect of LSD. This is suggestive of increased flexibility and novelty in brain dynamics induced by LSD compared with placebo, resembling the difference between improvisation (LSD) and memorized play (placebo) of a musical piece.

Taken together with previous studies associating psycho-pathology with deviations of brain dynamics from criticality^22,36,37,64^, this interpretation also suggests that the same dynamics that underlie creativity when tuned to criticality, may lead to psycho-pathology when the critical dynamics are impaired. Interestingly, this interpretation is supported by the shared genetic roots of schizophrenia, bipolar disorder, psychosis and creativity^72^, as well as by the shared network correlates of psychiatric disorders and creativity. For instance, the functional networks, whose activity and dynamic coupling is linked to creative thinking (DMN, SAL and ECN), show abnormal functional connectivity in psychiatric disorders such as schizophrenia^52,54,55^, bipolar disorders^56^, anxiety and depression^52^. Notably, discussions of psychological parallels between creativity and mental illness has a long history^73^.

Surprising parallels also emerge between the crucial role of RSN’s in psychiatric disorders and our multiple-correlation analysis relating the energy changes of different connectome harmonics to connectivity changes of the RSNs and to the intensity of different subjective experiences: In line with previous studies, highlighting the important role of abnormal DMN connectivity in psychiatric disorders such as depression^56,74^, anxiety^74^, bipolar disorder^56^ and schizophrenia^52,55^, as well as in the psychedelic state^10,61^, we found significant correlations between the DMN connectivity and the intensity of emotional arousal. This result suggests that the link between abnormal DMN connectivity and abnormal mental states may involve or even be mediated by altered emotional processing, also supporting previous studies linking DMN to emotional disregulation^57^.

Interestingly, all pairs of networks, whose connectivity changes showed significant correlation with the ratings of emotional arousal, i.e. DMN-SAL, DMN-rFP, DMN-pOP, PAR-pOP, have been previously reported to show increased between-RSN functional connectivity under LSD^17^, suggesting a potential link between this increased between-RSN functional coupling and emotional arousal experienced in the psychedelic state. In particular, abnormal DMN-SAL functional connectivity correlating with intense subjective effects has previously been reported under LSD^10^ and psilocybin^54^, and this coupling has been found to relate to ego-dissolution^9^, which is experienced as a positive feeling of oneness and loss of ego-boundaries. Our results also revealed significant correlations between the changes in coupled DMN-SAL connectivity and all three experiences of emotional arousal, positive mood and ego dissolution. This finding is also in agreement with and highly relevant for previous clinical^52,56^ and theoretical studies^52^ highlighting the important role of DMN-SAL coupling in various mood disorders such as depression, anxiety^52^, bipolar disorder^56^ and schizophrenia^52^.

In patients with schizophrenia, increased connectivity was also found between the lFP and temporal and parietal regions^75^. Although we acknowledge that chronic schizophrenia is a heterogeneous disorder (and not really a ‘state’ as such) with many phenomenological features that are inconsistent with the psychedelic state - and vice versa, notably our multiple correlation analysis revealed that the coupled connectivity changes of SAL with lFP alone or with FPN (lFP and rFP together) significantly correlated with the ratings of emotional arousal, positive mood and ego-dissolution, whereas connectivity changes of the SAL when coupled only with rPF did not yield the same level of significance. Moreover, the coupled connectivity of the lFP with the visual networks correlated with ratings of simple hallucinations, while the coupled connectivity of the rFP with the visual networks was found to significantly correlate with the ratings of both, complex imagery and simple hallucinations. These findings suggest that the cooperation between the visual networks and lFP and the lack of cooperation of the rFP may underlie related perceptual abnormalities seen not only in the psychedelic state but also in certain phases of psychosis, such as early psychosis^54^.

To conclude, here we have applied a new and powerful analytical methodology to the LSD state, yielding novel findings that inform not only on the neural correlates of this peculiar state of waking consciousness but also on the functioning of the brain more generally. The present findings highlight the value of viewing global brain function and related subjective states in terms of dynamic activation of harmonic brain states. Remarkably, this simple change in perspective reveals the dynamical repertoire of brain activity and suggests a shift of the brain dynamics towards whole-brain criticality under LSD. Importantly, by revealing the characteristic changes in cortical dynamics between LSD and normal awake state, the introduced method opens-up an opportunity for exploring the neural signatures of other psychological traits and states, including personality; creativity; psychiatric and neurological disorders; sleep, anaesthesia and disorders of consciousness; as well as other drug and non-drug induced altered states of consciousness.

## Methods

### Ethics statement

This study was approved by the UK National Health Service research ethics committee, West-London. Experiments were performed in accordance with the revised declaration of Helsinki (2000), the International Committee on Harmonization Good Clinical Practice guidelines, and National Health Service Research Governance Framework. Imperial College London sponsored the data collection, which was conducted under a Home Office license for research with schedule 1 drugs. All participants gave informed consent.

### Study design

Participants attended two sessions of scanning days (LSD and placebo) 14 days apart in a balanced-order, within-subjects design. The study was performed single-blind, where the staff were aware when LSD was being given but the participants were not. There were 2 non-music fMRI BOLD runs and one music fMRI BOLD run within each session. 70 minutes prior to MRI scanning each subject received either received received 75 *μ*g LSD in 10 mL saline (intravenous, I.V.) or placebo (10 mL saline, I.V.). Each BOLD scan lasted 7 min and the scans were completed 135 min postinfusion, as in^10^. The administration of LSD and placebo were counter-balanced such that half of the participants received LSD in scan one (Group 1) and half received it in scan 2 (Group 2) with a fixed and balanced assignment of participants into groups 1 and 2. To fully exclude any risk of direct carry over from the drug itself, the LSD and placebo sessions were separated by 14 days.

### Data

The fMRI blood oxygen level dependent (BOLD) scans were performed on 20 healthy subjects in 6 different conditions; LSD, placebo (PCB), LSD and PCB while listening to music, LSD and PCB after music session. Out of the 20 subjects, 12 were used for this analysis for the following reasons: one participant was excluded from analysis because of early termination of the scanning due to him reporting significant anxiety, 4 participants were excluded due to high levels of head movement (the criterion for exclusion for excessive head movement was subjects displaying higher than 15% scrubbed volumes when the scrubbing threshold is FD=0.5 as described in the original publication^10^), and 3 participants were excluded from the analysis due to technical problems with the sound delivery.

Participants were also asked to perform a limited number of visual analogue scale (VAS) style ratings at the end of each scan, using a button box in the scanner. Five key facets of the LSD experience were enquired about: 1) complex imagery (i.e. eyes-closed visions of objects, entities, landscapes etc.), 2) simple hallucinations (i.e. eyes-closed visions of shapes, colours, geometric patterns etc.), 3) emotional arousal (i.e. how emotional the participant felt, regardless of whether emotions were positive or negative), 4) positive mood, and 5) ego-dissolution (i.e. a fading sense of self, ego and/or subjectivity).

### Connectome harmonics as brain states

To characterize the spatio-temporal brain dynamics, we developed a new technique to decompose spatio-temporal recordings of brain activity into temporal evolution of brain states. The brain states are defined as spatial patterns formed by fully synchronized activity, each associated with a different spatial wavelength k and theoretically accompanying a different frequency of temporal oscillation in neural activity^15^. We estimate these fully synchronized brain states as the harmonics of macro-scale structural connectivity of the human brain - connectome harmonics - as described in^15^.

Recently, it has been shown that the harmonic modes of structural connectivity (human connectome), in fact predict patterns of correlated neural activity (functional connectivity)^15^. Furthermore, these harmonic modes of the human connectome, called *connectome harmonics*, yield a set of cortical patterns with increasing spatial frequency (indicated by wavenumber *k*) (Fig. S1). The distinct characteristic of the connectome harmonics is that they provide a spatial extension of the Fourier basis to the particular structural connectivity of the human brain. Thus, representing cortical activity as the combination of the activities of these brain states allows for a spatial frequency decomposition of any functional neuroimaging data. Notably, although intrinsically related (the higher the temporal frequency, the higher the wavenumber of the spatial pattern)^15^, the wavenumber of connectome harmonics that we investigate in this work should not be confused with the temporal frequency analysis performed by Fourier transform, but can be rather considered as its spatial extension to the particular structural connectivity of the human brain^15^.

### Computation of connectome harmonics

Connectome harmonics were estimated using an independent sample of participants, as also used in^15^, obtained and made available by the Human Connectome Project (HCP), WU-Minn Consortium (Principal Investigators: David Van Essen and Kamil Ugurbil; 1U54MH091657), which is funded by the 16 NIH Institutes and Centers that support the NIH Blueprint for Neuroscience Research and by the McDonnell Center for Systems Neuroscience at Washington University. We use magnetic resonance imaging (MRI) and DTI data of 10 unrelated subjects (six female, age 22-35) provided by the HCP, WU-Minn Consortium, available on https://db.humanconnectome.org. All MRI and DTI datasets were preprocessed according to minimal preprocessing guidelines of the HCP protocol and no additional preprocessing was performed.

To estimate the connectome-harmonic basis, we follow the exact same workflow as explained in^15^. For each subject, the cortical surfaces separating the white and grey matter were reconstructed from T1-weighted MRI data (resolution 0.7 mm), separately for each hemisphere using the Freesurfer Software http://freesurfer.net and represented as a graph with 10,242 nodes for each hemisphere. The white matter cortico-cortical and thalamo-cortical fibres were extracted from by applying a deterministic tractography algorithm^76^ using the MATLAB implementation of Vista Lab, Stanford University http://white.stanford.edu/newlm/index.php/MrDiffusion, to the diffusion tensor imaging (DTI) data of the same subjects. The DTI data and and the cortical surface were registered for each subject. Centred around each vertex (node)-in total 20,484-eight seeds were initialized and the tractography was performed with the following parameters: fractional anisotropy (FA) threshold 0.3, i.e. FA<0.3 being termination criteria for the tracking, minimum tract length 20 mm, and maximum angle between two contiguous tracking steps 30 degrees.

After forming a graph representation of the human connectome 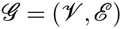, where the vertices were sampled form the surface of gray matter by the nodes 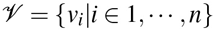 with *n* being the total number of nodes and the edges represent the local as well as the long-range connections (estimated via tractography) between the vertices 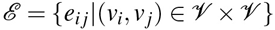 we use an undirected, unweighted graph to represent the adjacency (connectivity) matrix of each subject:

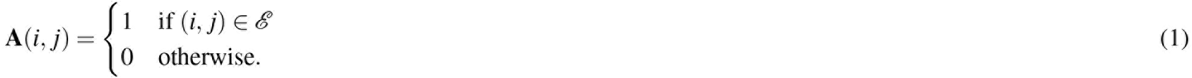

Finally we average the adjacency matrices of 10 subjects yielding a group average adjacency matrix **Ā** to represent the average structural connectivity of all subjects. Again following the study in^15^, we compute the symmetric graph Laplacian 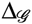 on the average connectome graph (average adjacency matrix) in order to estimate the discrete counterpart of the Laplace operator^77^ Δ applied to the human connectome; i.e. the connectome Laplacian, as:

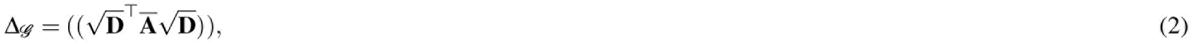

where the adjacency matrix **A** is defined in Eq. (1) and

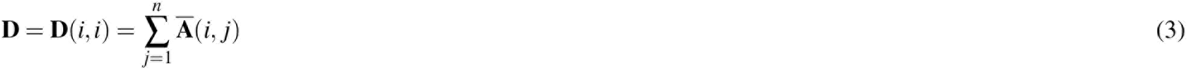

denotes the degree matrix of the graph. We then calculate the connectome harmonics *ψ*_*k*_, *k* ∈ {1, ···, *n*} by solving the the following eigenvalue problem:

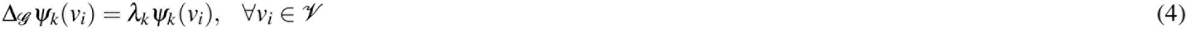

with λ_k_, *k* ∈ {1, ···, *n*} being the corresponding eigenvalues of 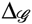.

### Connectome-harmonic decomposition of fMRI data

To represent the fMRI data acquired in each different condition in the cortical surface coordinates of connectome harmonics, each fMRI scan was projected onto the cortical coordinates using the *-volume-to-surface-mapping* command of the Human Connectome Project Workbench. This registration yields the time course 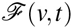 for all vertices 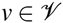 on the cortex. Then, the spatial cortical activity pattern 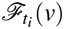 for each time point *t*_*i*_ ∈ [1, …, *T*] of the fMRI data 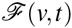 was decomposed into the activity of connectome harmonics 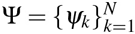

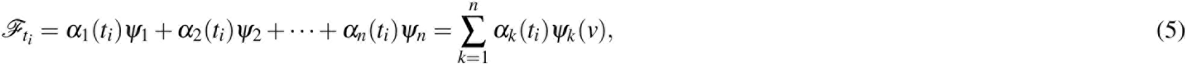

where the temporal activity *α*_*k*_(*t*) of each connectome harmonic *ψ*_*k*_ was estimated by projecting the fMRI data 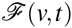 onto that particular harmonic: *α*_*k*_ are estimated as:

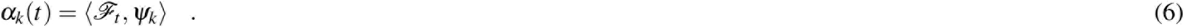

Note that the subscripts *k* and *t* denote the wavenumber of the corresponding connectome harmonic *ψ*_*k*_ and time instance, respectively.

### Power and energy of brain states

Power of each connectome harmonic *ψ*_*k*_, *k* ∈ [1, ···, *n*] in the cortical activity pattern at a particular time point *t* in the fMRI data is computed as the strength of activation of a particular connectome harmonics *ψ*_*k*_:

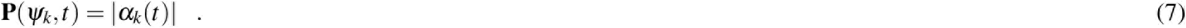

Energy of each connectome harmonic *ψ*_*k*_, *k* ∈ [1, ···, *n*] in the cortical activity pattern at a particular time point *t* in the fMRI data is estimated by combining the strength of activation of a particular connectome harmonics with its own intrinsic energy given by 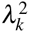 (Eq. 4). Hence, we define the energy of a brain state *ψ*_*k*_ as:

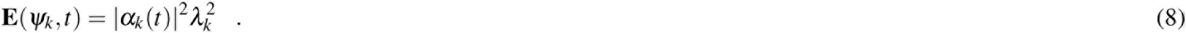

The total energy of brain activity for any given time point *t* is then given by:

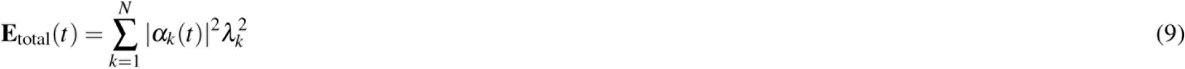

which also is equal to

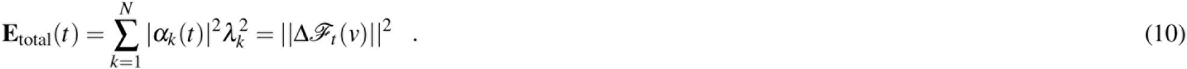

Note that Laplace operator Δ measures the amount of flow of activity and hence the total energy of brain activity corresponds to the total amount of flow of neural activity throughout the cortex at a particular time point *t*. The total energy and power of a brain state are computed by summing over all time points. Considering that the power is defined as the dot product between the pattern of cortical activity at a given time and a connectome harmonic, and given that the connectome harmonics are orthonormal; i.e. ‖ *ψ*_*k*_‖ = 1, ∀*k*, the upper bound of the power is determined by the cortical activity pattern, whereas the lower-bound is 0. In case of the energy values the square of power is weighted by the square of the corresponding eigenvalue of the connectome harmonic. The eigenvalues are bounded by *λ*_*k*_ < [0, ···, 2] for the connectome Laplacian used in this study^77^.

### Power-law analysis

When plotted on a log_10_-log_10_ plot, power-laws follow a straight line with a slope equal to their critical exponent *β*^37^. We evaluate the relations between maximum power, average power as well as power fluctuations and the wavenumber *k* of over the whole spectrum of connectome harmonics in log_10_ coordinates. Logarithmic binning with 100 bins is performed to smooth the curves^40^. The line fitting and the estimation of the critical exponent *β* was performed in MATLAB. Goodness of fit of the line in log_10_-log_10_ space was measured as the root mean square error *ε* of the line fit. For all comparisons using t-tests, the power-laws were estimated separately for each subject and each conditions and the t-tests between LSD and PCB conditions were performed on the distributions of the root mean square error ε and critical exponent *β* across all subjects.

### Cross-frequency correlations between brain states

Cross frequency correlations between all pairs brain states (*ψ*_*i*_, *ψ*_*j*_) are estimated as:

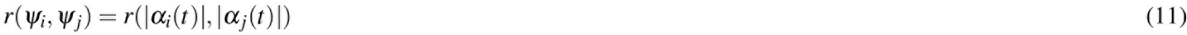

where *r* denotes Pearson’s linear correlation coefficient.

### Estimation of RSN connectivity

RSNs were estimated using an independent sample of participants, as also used in^10^, as part of the Human Connectome Project (HCP), WU-Minn Consortium (Principal Investigators: David Van Essen and Kamil Ugurbil; 1U54MH091657). Estimation of the RSNs and RSN connectivity was performed as described in the original publication^10^.

### Multiple correlations between RSN connectivity and subjective ratings

Multiple correlations between RSN connectivity and subjective ratings were estimated using multiple correlation coefficient^53^. First, for each subject, the functional connectivity changes within each RSN were estimated between the LSD and PCB conditions for the scans without music. For each RSN, combining the functional connectivity (FC) changes across all 12 subjects and 2 scans results in a 24 dimensional vector. In the direct application of multiple correlation coefficient^53^, correlations were evaluated directly between these 24 dimensional vectors representing RSN connectivity changes and the subjective ratings of the corresponding scans (again represented as 24 dimensional vectors (12 subjects × 2 scans)). In the indirect application of the multiple correlation coefficient^53^, each 24 dimensional vector of RSN connectivity and subjective rating is first correlated with the changes in energy of a set of connectome harmonics between LSD and PCB conditions comparable to a spectral decomposition of the 24 dimensional vectors. Multiple correlations between RSN connectivity changes and subjective ratings were than evaluated on the connectome-harmonic correlates of the 24 dimensional vectors. Fig. S4 illustrates the evaluation of different number of connectome-harmonic correlations used to represent the 24 dimensional vectors of FC changes and subjective ratings. This allows for mapping the data (24 dimensional vectors of FC changes and subjective ratings) into a higher dimensional space, where the maximum dimensionality is equal to the number of brain states and hence representing the same information with a higher resolution.

### Data availability

The datasets analysed during the current study can be made available on request.

## Acknowledgements

The authors would like to thank Andrea Insabato for valuable discussions on significance analysis.

## Author contributions statement

R.L.C.H. designed and managed experimental data collection. L.R. and M.K. assisted the data collection and performed the pre-processing. S.A. designed and carried out the analyses and statistical testing, with contributions from G.D. and M.L.K., S.A., G.D., M.L.K and R.L.C.H. wrote the manuscript together.

